# Aggressive but not reproductive boldness in male green anole lizards correlates with baseline vasopressin activity

**DOI:** 10.1101/2021.09.11.459908

**Authors:** David Kabelik, Allison R. Julien, Brandon R. Waddell, Mitchell A. Batschelett, Lauren A. O’Connell

## Abstract

Across species, individuals within a population differ in their level of boldness in social encounters with conspecifics. This boldness phenotype is often stable across both time and social context (e.g., reproductive versus agonistic encounters). Various neural and hormonal mechanisms have been suggested as underlying these stable phenotypic differences, which are often also described as syndromes, personalities, and coping styles. Most studies examining the neuroendocrine mechanisms associated with boldness examine subjects after they have engaged in a social interaction, whereas baseline neural activity that may predispose behavioral variation is understudied. The present study tests the hypotheses that physical characteristics, steroid hormone levels, and baseline variation in Ile^3^-vasopressin (VP, a.k.a., Arg^8^-vasotocin) signaling predispose boldness during social encounters. Boldness in agonistic and reproductive contexts was extensively quantified in male green anole lizards (*Anolis carolinensis*), an established research organism for social behavior research that provides a crucial comparison group to investigations of birds and mammals. We found high stability of boldness across time, and between agonistic and reproductive contexts. Next, immunofluorescence was used to colocalize VP neurons with phosphorylated ribosomal protein S6 (pS6), a proxy marker of neural activity. Vasopressin-pS6 colocalization within the paraventricular and supraoptic nuclei of the hypothalamus was inversely correlated with boldness of aggressive behaviors, but not of reproductive behaviors. Our findings suggest that baseline vasopressin release, rather than solely context-dependent release, plays a role in predisposing individuals toward stable levels of displayed aggression toward conspecifics by inhibiting behavioral output in these contexts.

## Introduction

Humans are often described as being introverted, extraverted, or somewhere in between. Such variation in a stable boldness measure similarly exists within numerous animal species, and has also been described as a behavioral syndrome, personality, or coping style (Koolhaas et al., 2010; Réale et al., 2010; Sih et al., 2004). These terms encapsulate the notion of linked social traits that are expressed across various environments, including both social and nonsocial contexts. Indeed, evidence in some species demonstrates heritability of displayed boldness (Ballew et al., 2017; Mont et al., 2018; Scherer et al., 2017), suggesting a genetic basis for the stability of individual differences across time. Likewise, the level of an individual’s boldness is often consistent across social contexts (Colléter and Brown, 2011; Koolhaas et al., 2010; Qu et al., 2018; Reaney and Backwell, 2007), where individuals that may be shy within a reproductive social encounter may also be shy within an agonistic encounter (Kabelik et al., 2021). Such consistency of behavioral phenotype hints at common neural mechanisms regulating boldness across a variety of contexts.

The mechanisms that underlie boldness during social encounters likely involve the social decision-making network (Newman, 1999; O’Connell and Hofmann, 2012, 2011) and its various neuroendocrine mediators (Baugh et al., 2012; Félix et al., 2020; Ketterson and Nolan Val, 1999) that are conserved across vertebrates. The mediators linked to individual variability in behavioral phenotype include steroid hormones (Koolhaas et al., 2010; Sluyter et al., 1996; Tudorache et al., 2018; Veenema et al., 2004, 2003), various neuropeptide and neurotransmitter systems (Thörnqvist et al., 2019; Veenema et al., 2004), and their associated receptors (Alfonso et al., 2019; Kabelik et al., 2021; Kanitz et al., 2019). Although much of the research on the neuroendocrine basis of boldness in social contexts has been conducted in mammals and fish, here we examine these traits in a reptilian model system, the green anole lizard (*Anolis carolinensis*). The study of behavioral neuroscience in reptiles lags behind that of other vertebrate groups and invertebrates (Kabelik and Hofmann, 2018; Taborsky et al., 2015), despite lizards being important for evolutionary comparisons among animal taxa, especially amniotic vertebrates (mammals, birds, and reptiles). Green anoles are a longstanding model for behavioral neuroendocrinology research (Lovern et al., 2004) due to their readily quantifiable social behavioral displays, natural seasonal variability in hormone levels, and natural stress responsivity. A social decision-making network has been described in reptiles (Kabelik et al., 2018), and boldness has been shown to be stable across contexts and over time in anole lizard studies (Kabelik et al., 2021; Putman et al., 2019).

Various neuroendocrine variables have been related to the expression of social behaviors in lizards, including neuropeptides and sex steroid hormones (Dunham and Wilczynski, 2014; Hartline et al., 2017; Kabelik et al., 2013, 2008b; Kabelik and Crews, 2017; Kabelik and Magruder, 2014; Korzan et al., 2001; Korzan and Summers, 2004; Larson and Summers, 2001; Smith and Kabelik, 2017; Watt et al., 2007; Woolley et al., 2004a, 2004b, 2001). Here, we focus on the potential regulation of boldness by the vasopressin (VP; Ile^3^- vasopressin, a.k.a. Arg^8^-vasotocin) system, as well as circulating steroid hormones. Vasopressin has been shown to have various effects on social behavioral expression, and stress reactivity, across species (Albers, 2015; Carter, 2017; Goodson and Kabelik, 2009; Kelly et al., 2011; Kelly and Goodson, 2014a; Walton et al., 2010; Wilczynski et al., 2017). Depending where in the brain VP is released, the effects on social behavior may be completely opposed to each other, such as in the case of displayed aggression in rats (Veenema et al., 2010). Research in anole lizards suggests that VP has an inhibitory effect on components of the social decision-making network (Kabelik et al., 2018). Although many studies focus on activity of VP neurons and receptors during a social encounter (e.g., Kabelik et al., 2013), information about baseline activity of VP neurons in individuals varying in boldness is unknown.

Here, we test three hypotheses about the regulation of boldness with social contexts of male green anoles. First, we test the hypothesis that the physical size of males relates to their boldness during social encounters. Body size has been shown to be predictive of boldness in some but not all species (Adriaenssens and Johnsson, 2011; Brown and Braithwaite, 2004; Mayer et al., 2016; Smith and Blumstein, 2010), and thus we predicted either a positive association or no association between these variables. Second, we test the hypothesis that circulating levels of steroid hormones regulate boldness within social behavior interactions. As testosterone and progesterone have been linked to aggression (Kabelik et al., 2008b; Weiss and Moore, 2004) and glucocorticoids to boldness in other species (Koolhaas et al., 2010; Sluyter et al., 1996), we predicted that steroid hormone levels would predict behavioral boldness in the present study. Third, we tested the hypothesis that central release of VP modulates boldness within social interactions. As VP is associated with both affiliative and agonistic behavior (Kelly and Goodson, 2014b), we predicted a relationship between these variables, but not predict a specific direction of this correlation.

We first examined the stability of boldness in both reproductive and agonistic encounters across a period of two weeks. We then waited for one day before collecting blood samples and brains to eliminate neuroendocrine changes correlated with recent participation in a social behavior interaction. We quantified hormone levels via enzyme-linked immunosorbent assay and used immunofluorescence to label VP neurons and their colocalization with phosphorylated ribosomal protein S6 (pS6), a proxy marker of neural activation in response to stimulation, as well as baseline neural activity (Cao et al., 2011; Klingebiel et al., 2017; Knight et al., 2012). We predicted that circulating steroid hormone levels and the baseline activity of VP neurons (%VP-pS6 colocalization) would reflect the neuroendocrine state that predisposes that individual toward boldness or shyness.

## Materials and Methods

### Subjects

Twenty-two male green anoles (*Anolis carolinensis*) were obtained from a commercial supplier and served as our focal subjects. They were singly housed for three weeks prior to experimentation within terraria (30.5 cm H x 26 cm W x 51 cm L) and kept in breeding season conditions: long-day (14 light:10 dark) full-spectrum lighting, 12 hours of supplemental heat provided 5 cm above one end of a wire-mesh terrarium lid by means of a 60-W incandescent light bulb. Animals were fed three times per week with crickets and cages were misted twice-daily with deionized water. Additional males and females from our housing colony were used as stimulus animals in social interactions. All procedures involving live animals were conducted according to federal regulations and approved by the Institutional Animal Care and Use Committee at Rhodes College.

### Behavior assays

Behavioral trials were carried out in May and June of 2013. Focal males were assessed for boldness within two social encounter scenarios – a reproductive encounter with two adult conspecific females and an agonistic encounter with a size-matched (within 3 mm snout-vent length) conspecific stimulus male. In each case, the stimulus animals were placed into the focal male’s terrarium and behaviors were scored for 10 minutes. Two females were used in the reproductive encounter to maximize the probability of eliciting reproductive behaviors from the focal male. We recorded the frequency (sum of behaviors per 10-min session) and latency to first performance (minute of first occurrence of any listed behavior) of the following behaviors: head bob bout, push-up bout, dewlap extension bout, dewlap extension bout with push up, chase, and copulate. Focal males that failed to display any behaviors were assigned the maximum latency score of 10 min. The maximum intensity of behavioral display was also scored from 0-4 based on the highest achieved category: no display, headbob only, pushup and/or dewlap display, chase, and copulate. Behaviors in the agonistic encounter were scored as in the reproductive encounter, except that biting of the stimulus male replaced copulation as the highest intensity behavior.

The reproductive encounter was conducted with three separate pairs of females and the agonistic encounter with three separate stimulus males over the course of a week. Only one trial was conducted per day. Stimulus animals were only used once per day. The entire procedure was then repeated during the subsequent week, toward the same three pairs of females and the same three stimulus males. Conducting the behavioral observations a second time allowed us to determine the stability of each boldness measure across time.

### Bold-shy categorization

To generate each boldness score, we conducted principal components analyses (PCA) to reduce the average behavioral latency, frequency, and intensity scores from the given encounter scenarios (reproductive or agonistic), separately for week 1 and for week 2, each into a single value. For example, in male-female trials from week 1, the behavioral latency, frequency, and intensity scores for each focal male were averaged across the three trials to generate an “average reproductive latency”, “average reproductive frequency”, and “average reproductive intensity” score. These average scores were then included in the PCA and generated a single PCA component that we here call BoldnessToFemalesWeek1. In each scenario, the resulting analysis generated a single PCA component with an eigenvalue > 1, and in each case, this component was highly positively correlated with average frequency and intensity scores, and negatively with average latency scores (r > ± 0.80, p < 0.001 for each). BoldnessToFemalesWeek1 explained 81% of the behavioral variation in those trials, BoldnessToFemalesWeek2 (from week 2 data) explained 75% of the variation from those data, BoldnessToMalesWeek1 (from week 1 agonistic encounters) explained 83% of the variation in those data, and BoldnessToMalesWeek2 explained 85% of the variation in behavioral frequency, latency, and intensity values from the week 2 agonistic encounter data. As week 1 and week 2 data were highly correlated, we also created a variable that averaged reproductive boldness from weeks 1 and 2 (AverageBoldnessToFemales), as well as agonistic boldness from weeks 1 and 2 (AverageBoldnessToMales).

### Tissue harvesting

Prior to handling for blood and brain harvesting, focal subjects were left undisturbed in their home terraria for 1 day following their last behavioral trial. We euthanized focal males by cutting through the spinal column and immediately collected trunk blood for hormone analyses (average collection time from first handling was 162 ± 3.2 s). The blood was kept at 4°C until centrifugation. The brain was then rapidly dissected and fixed by overnight submersion in 4% paraformaldehyde in 0.1 M phosphate buffer at 4°C, followed by cryoprotection with 30% sucrose in 0.1 M phosphate-buffered saline (PBS). The body (minus the head) was then weighed, after which the testes were dissected from the body and weighed. Brains were sectioned into two series, at a section thickness of 50 µm, on a Microm HM 520 cryostat (Thermo Scientific).

### Hormone analyses

Blood samples were centrifuged and plasma (averaging 62 ± 2.7 µl) was frozen at -80°C until hormone analysis. We quantified testosterone (ADI-900-065; sensitivity 5.67 pg/mL), estradiol (ADI-900-008; sensitivity 28.5 pg/mL), progesterone (ADI-900-011; sensitivity 8.57 pg/mL), and cortisol (ADI-900-071; sensitivity 56.72 pg/mL) using enzyme-linked immunosorbent assay (ELISA) kits (Enzo Life Sciences, Farmingdale, NY). The cortisol kit cross-reacts with corticosterone at 28%, representing a general glucocorticoid assay, albeit with lower-than-typical sensitivity. We re-suspended 7 µl of plasma in 203 µL of the appropriate assay buffer and ran each sample in duplicate as per manufacturer’s instructions. Samples were run across two plates for each hormone and the inter-assay variation across plates and the intra-assay variance for each plate is as follows: testosterone (inter: 5.6%; intra: 5.6% and 6.1%), estradiol (inter: 4.1%; intra: 3.7% and 8.5%), progesterone (inter: 4.7%; intra: 2.3% and 4.7%), and cortisol (inter: 6.4%; intra: 3.8% for both plates). Five samples were inadvertently excluded from the analysis. Hormone results were generally consistent with previously reported levels in this species (Greenberg and Crews, 1990; Young et al., 1991).

### Immunohistochemistry

Immunohistochemical processing was conducted as in previous studies (Hartline et al., 2017; Kabelik et al., 2018, 2014, 2013; Kabelik and Magruder, 2014). Briefly, we processed one series of brain sections with 1:250 dilution of rabbit anti-pS6 antibody (#2211, Cell Signaling Technology), 1:5000 dilution of guinea pig anti-vasopressin antibody (T-5048, Peninsula Laboratories), and 1:2000 dilution of sheep anti-tyrosine hydroxylase antibody (NB300-110, Novus Biologicals; part of a separate study). The sections were subsequently processed with donkey anti-sheep secondary antibody conjugated to Alexa Fluor 488 at 3 µl/ml and (Life Technologies), donkey-anti rabbit secondary antibody conjugated to Alexa Fluor 555 at 5 µl/ml (Life Technologies), and donkey-anti guinea pig DyLight 647 at 16 µl/ml (Jackson ImmunoResearch). Preadsorption with excess VP (4100576, Bachem) and pS6 blocking peptide (1220S, Cell Signaling Technology) antibody eliminated signal (**Supplementary Fig. 1**). We targeted pS6 rather than Fos because preliminary research demonstrated extremely low levels of Fos expression within VP neurons under baseline conditions.

### Microscopy and Image Analyses

An LSM 700 Confocal microscope and Zen 2010 software (Carl Zeiss), using a 20X objective, were used to capture z-stacks of photomicrographs at 5 µm intervals, in a grid that was later stitched together. A maximum intensity projection created a two-dimensional image. Individual colors were exported as separate layers using AxioVision 4.8 (Carl Zeiss), and these were stacked as overlaid monochromatic layers in Photoshop (Adobe Systems). Layers in the stack could thus be toggled on and off to determine signal colocalization. Analyses were conducted by individuals unaware of treatment groups.

We examined VP cells within the paraventricular nucleus (PVN) and the supraoptic nucleus (SON) of the hypothalamus (**Fig. 1**). Only cells that could be clearly visualized with VP-signal in the cytoplasm and a darker nucleus were examined. VP cell counts could not be accurately obtained due to damage to some tissue sections and many overlapping cells, especially within the SON. We estimated VP cell density by examining the average number of cells per section across the three most densely populated sections. An average of 39.32 ± 4.93 (mean ± S.E.) cells in the PVN and 28 ± 5.85 cells in the SON were analyzed per subject.

**Figure 1.**
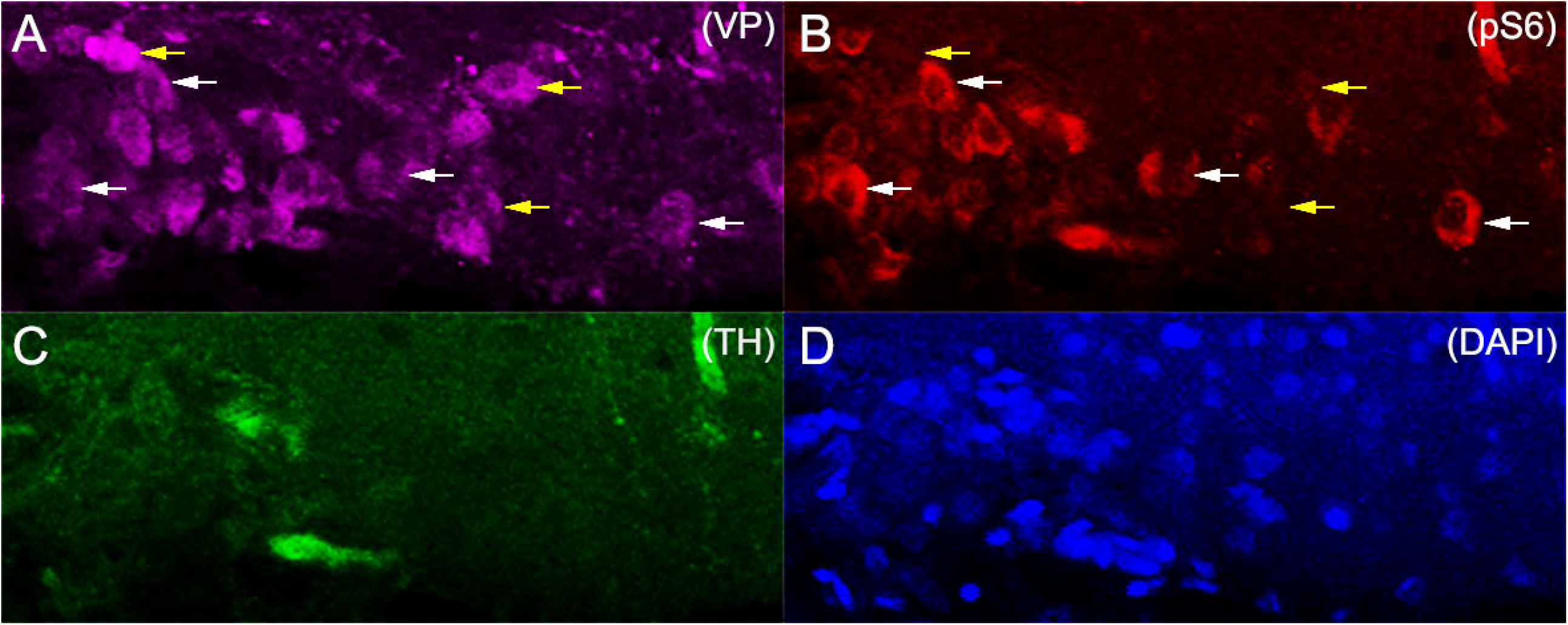
Immunofluorescence within the SON. VP-immunoreactive cells (A) were scored as colocalized (white arrows) or not colocalized (yellow arrows) with pS6-immunoreactive signal (B). Nonspecific signal due to the presence of blood cells was disregarded because those cells would also cause autofluorescence within the tyrosine hydroxylase (TH) layer (C; captured for a separate study). A DAPI-stained layer was also captured (D).

### Statistical analyses

Statistical analyses (Pearson’s correlations; repeated-measures analysis of variance, ANOVA; Friedman test; PCA) were run using IBM SPSS (version 22). Corrections for multiple comparisons were made using Benjamini-Hochberg calculations (Benjamini and Hochberg, 1995). Scatterplots and boxplots were made with ggplot2 (version 3.3.3) in RStudio (version 1.4.1106) running R (version 4.1.0). Hormone levels were ln-transformed to meet assumptions of parametric analyses.

## Results

### Boldness is stable over time and correlated across contexts

Relative boldness was found to be stable across weeks (**Fig. 2**). BoldnessToMalesWeek1 was highly correlated with BoldnessToMalesWeek2 (r=0.57, N=22, p=0.005). This was despite a general drop in aggression frequency and intensity, and rise in aggression latency across the six testing sessions, possibly due to habituation (p<0.01 for all three variables, see **Supplementary Figs. 2-4**). Similarly, BoldnessToFemalesWeek1 was highly correlated with BoldnessToFemalesWeek2 (r=0.72, N=22, p<0.001). Reproductive behavior frequency, intensity, and latency did not differ across testing sessions (p>0.05 for all, see **Supplementary Figs. 5-7**). Because the week 1 and week 2 boldness PCA scores were highly correlated, the average scores from both weeks were then used to compare boldness across social contexts, where AverageBoldnessToFemales correlated strongly with AverageBoldnessToMales (r=0.65, N=22, p=0.001).

**Figure 2.**
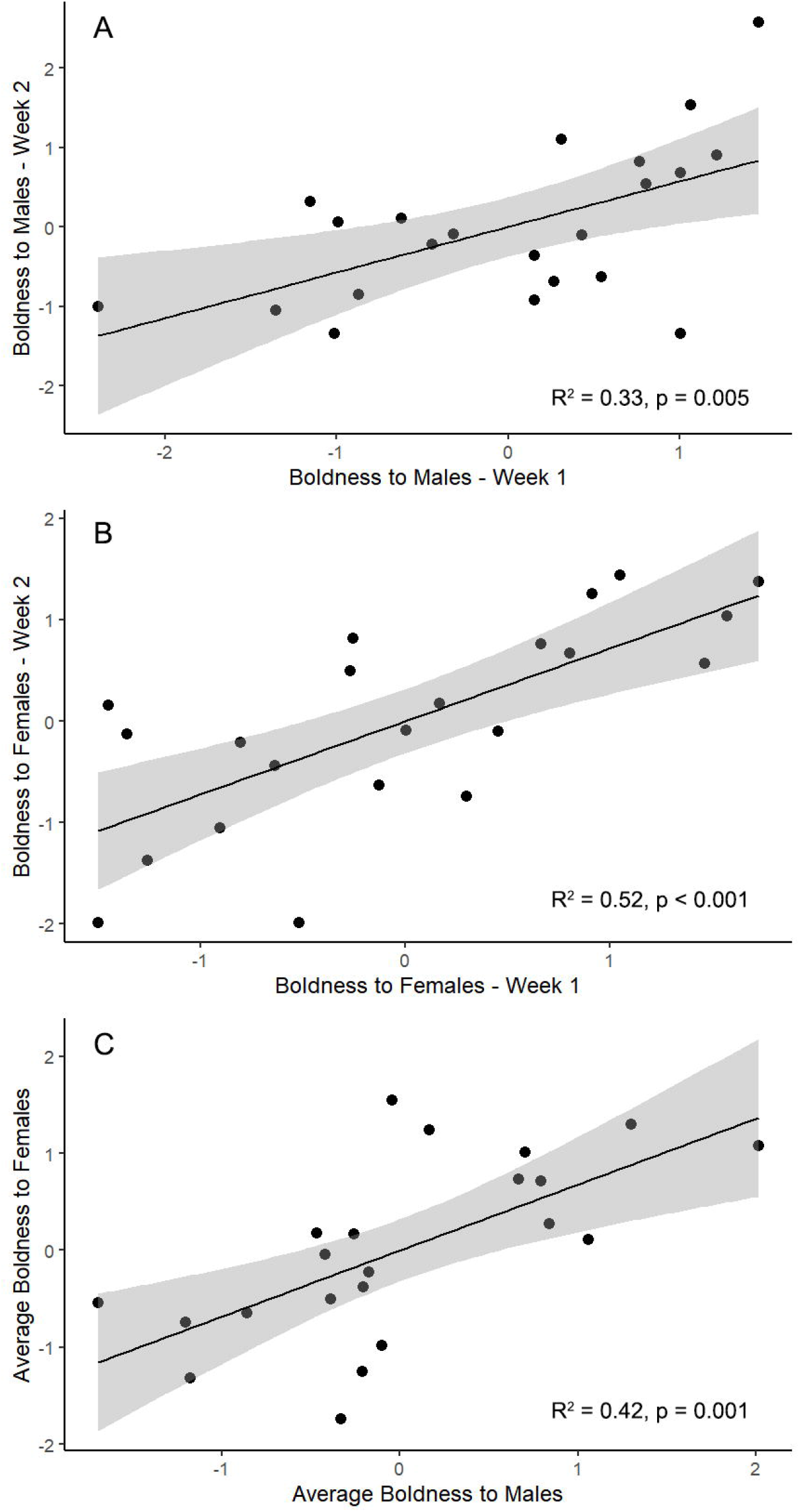
Stability of boldness. Boldness scores were correlated across weeks within agonistic contexts involving encounters with three separate conspecific intruder males across two separate weeks (A), as well as within reproductive contexts involving encounters with three separate pairs of conspecific females across separate weeks (B). Average boldness scores across the two weeks in the agonistic context were highly correlated with average boldness score within the reproductive context (C).

### Boldness is generally not correlated with physical traits or steroid hormone levels

Boldness to males and females was unrelated to physical or hormonal measures except for a positive correlation between testes mass and AverageBoldnessToFemales (**Table 1**). Similarly, neither VP-pS6 colocalization within the PVN, nor in the SON, correlated with any physical characteristics or hormone levels (p>0.05 for all).

**Table 1.** Results of correlations between measures of boldness and physical and hormonal measures. Displayed are the Pearson correlation coefficient (r), the sample size (N), and the probability of significance (P). No correlations were significant following correction for multiple comparisons.

|  | AverageBoldnessToMales |  |  | AverageBoldnessToFemales |  |  |
| --- | --- | --- | --- | --- | --- | --- |
|  | r | N | P | r | N | P |
| **Physical Variables:** |  |  |  |  |  |  |
| snout-vent length (mm) | -0.02 | 22 | 0.94 | -0.13 | 22 | 0.55 |
| body-minus-head mass (g) | -0.10 | 22 | 0.67 | -0.10 | 22 | 0.67 |
| testes mass (g) | 0.31 | 22 | 0.16 | 0.47 | 22 | 0.026 |
| testosterone (ng/ml) | 0.21 | 17 | 0.42 | 0.29 | 17 | 0.25 |
| estradiol (ng/ml) | 0.12 | 17 | 0.65 | 0.31 | 17 | 0.22 |
| progesterone (ng/ml) | 0.12 | 17 | 0.65 | 0.28 | 17 | 0.28 |
| glucocorticoids (ng/ml) | -0.20 | 17 | 0.45 | 0.17 | 17 | 0.51 |

### Boldness to males was associated with vasopressin activity in the PVN

AverageBoldnessToMales was negatively correlated with VP-pS6 colocalization (**Fig. 3**) within the PVN (r=-0.51, N=22, p=0.014) and the SON (r=-0.52, N=22, p=0.014). AverageBoldnessToFemales was not correlated with VP-pS6 colocalization in either the PVN (r=-0.24, N=22, p=0.29) or the SON (r=-0.09, N=22, p=0.69).

**Figure 3.**
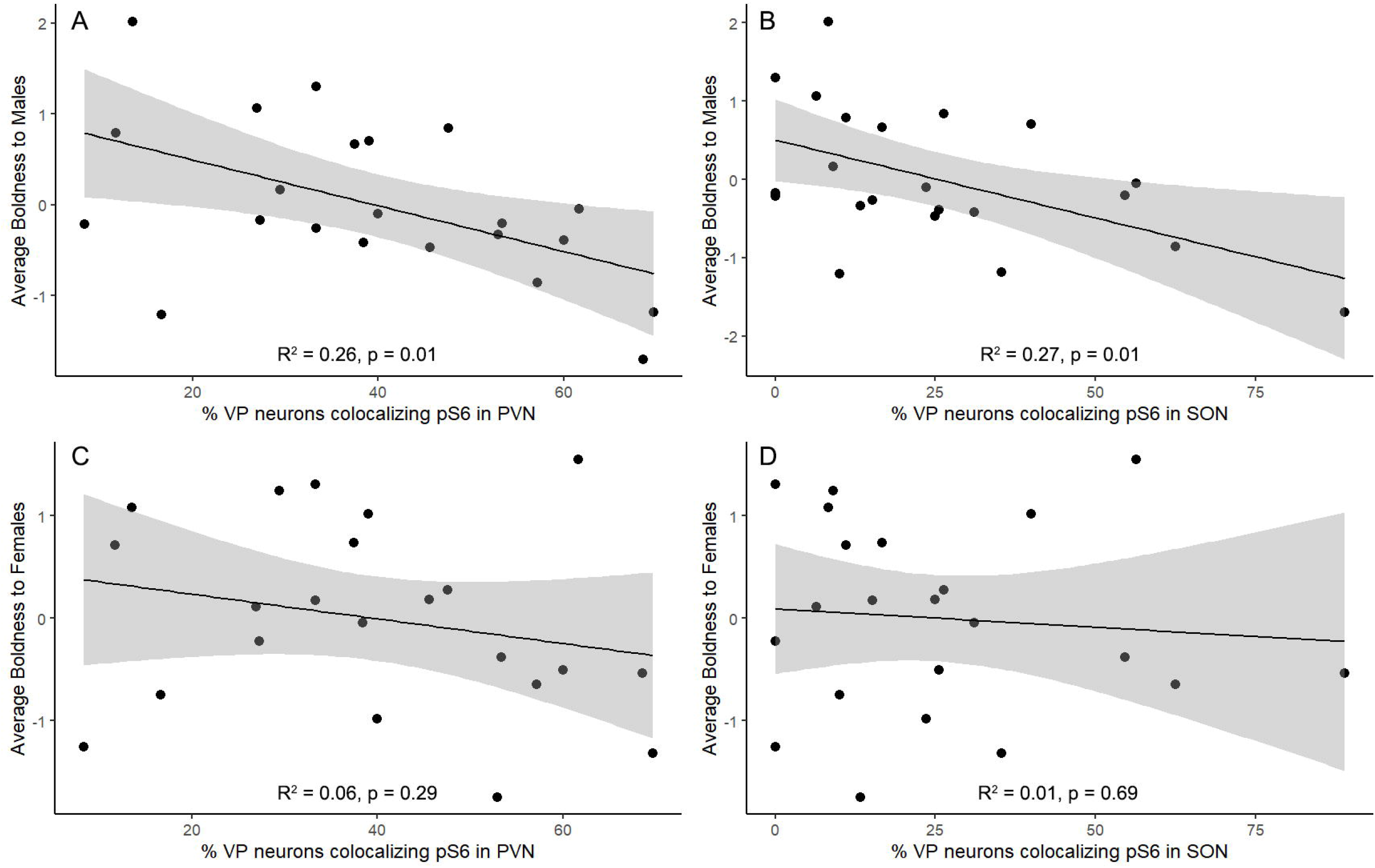
Boldness of social behavior interactions relative to VP-pS6 colocalization. Average boldness scores toward males within an agonistic context (A, B) but not females within a reproductive context (C, D) correlated negatively with the percentage of VP neurons in the PVN and SON that were pS6 positive.

### Boldness to males and females was not associated with measures of total VP cell number

The average PVN VP cell density across the three most densely populated sections was unrelated to displayed boldness to males (r=-0.05, N=22, p=0.84) and females (r=0.22, N=22, p=0.33). This was likewise true for SON VP cell density and displayed boldness to males (r=-0.19, N=20, p=0.42) and females (r=-0.02, N=20, p=0.95). Similar analyses for average VP counts per section, or VP counts on the single most densely populated section likewise showed no relationship to displayed boldness (p>0.27 for all).

## Discussion

In this study, we tested the hypotheses that physical size, circulating steroid hormone levels, and central release of VP regulate displayed boldness in reproductive and agonistic social encounters. Our results support the third hypothesis, where baseline levels of VP activity are associated with boldness of male green anoles within an agonistic context, presumably due to differential levels of VP release. However, our results do not support our first two hypotheses relating to physical size and steroid hormones levels being associated with boldness. We also demonstrate that boldness in male green anoles is stable across social contexts and time.

Vasopressin is a neuromodulator that has previously been shown to have causal effects on aggression (Kelly and Goodson, 2014b; Terranova et al., 2017), and our study supports the notion that basal VP activity helps to determine individual differences in aggressive behavior output. Our results are in line with previous findings showing a negative correlation between neuronal VP activity and activation of social decision-making network nodes in the closely related brown anole, especially within agonistic contexts (Kabelik et al., 2018). Interestingly, although boldness measures are correlated between agonistic and reproductive contexts, VP activity does not correlate well with reproductive boldness suggesting that other mechanisms may underlie the cross-context correlation.

### Stability of boldness

Our results demonstrate stability of displayed boldness across weeks and social contexts. The correlation of boldness scores across contexts suggests a shared neural network that regulates general social behavioral output. Correlated boldness measures are often referred to as behavioral syndromes (Colléter and Brown, 2011; Koolhaas et al., 2010; Qu et al., 2018; Reaney and Backwell, 2007). The stability of boldness across time, on the other hand, supports the notion that these traits are at least partly hard-wired, as would be expected if boldness has strong heritable components, as has been suggested by previous studies (Ballew et al., 2017; Mont et al., 2018; Scherer et al., 2017). Various selective pressures appear to maintain variability in exhibited boldness within populations (Koolhaas et al., 2010; Smith and Blumstein, 2010).

### Boldness is unrelated to physical characteristics

Although a relationship between body size and boldness has been demonstrated in reptiles, this relationship was observed in juvenile keelback snakes emerging from shelter (e.g., Mayer et al., 2016), and we did not find any such relationships in the present study of adult green anoles. Instead, our results were very much in line with those of Kabelik et al. (2021), where boldness of social displays was not found to correlate with measures of body size. This was true for both boldness within agonistic and reproductive contexts. The one physical variable that did show a correlation trend with boldness was testes mass. However, this result is inconsistent with the findings of Kabelik et al. (2021) that included a larger sample size and did not find any such relationship. Moreover, the present finding did not survive correction for multiple comparisons.

### Boldness is unrelated to circulating steroid hormone levels

We predicted that glucocorticoids would correlate with boldness in green anole lizards based on studies that found differences in glucocorticoid levels between individuals differing in active versus passive coping styles (Koolhaas et al., 2010; Sluyter et al., 1996). However, as in Kabelik et al. (2021), we found no relationship between circulating glucocorticoid levels and displayed boldness during social interactions in the present green anole study. It is nevertheless important to note that our hormone measures were from baseline diurnal plasma samples, and thus we cannot exclude the possibility that nocturnal or stress-evoked glucocorticoid levels may still relate to exhibited boldness. This is especially true given the fact that VP is a releasing hormone for adrenocorticotropic hormone, including in birds (Cornett et al., 2013), thus suggesting a similarly conserved role in reptiles.

We originally also hypothesized that sex steroid hormones may influence boldness in male green anoles because previous lizard studies demonstrated their involvement in the regulation of social behaviors. For instance, in male tree lizards, circulating testosterone correlates with aggression (Kabelik et al., 2006) and testosterone and progesterone treatments causally promote aggression (Kabelik et al., 2008b; Weiss and Moore, 2004). Additionally, in male brown anoles, the anti-androgen cyproterone acetate reduced display behaviors toward conspecific males and females (Tokarz, 1995). In male side-blotched lizards, the more aggressive morph type has also been shown to possess higher levels of testosterone (Sinervo et al., 2000). However, in the present study, we did not find any correlations between baseline sex steroid hormone levels and measures of displayed boldness within social contexts, which is in line with Kabelik et al. (2021). However, that study did detect differences in androgen receptor expression in the ventromedial hypothalamus between bold and shy males, suggesting that activational androgen signaling may nevertheless be involved in regulating the boldness of social behaviors. Furthermore, organizational sex steroid levels likely also play a role in determining levels of adult boldness.

### Building upon previous VP research in reptiles

Previous research examining VP cell numbers and optical densities in tree lizards found no correlations with aggression frequency or intensity (Kabelik et al., 2008c). However, other lizard studies provided findings suggestive of an involvement of VP in the regulation of social behaviors. For instance, neural VP expression was found to be higher in dominant green anoles than in subordinate animals (Hattori and Wilczynski, 2009). Furthermore, rather than solely examining VP expression, a brown anole study examining the colocalization of VP neurons with Fos (another measure of neural activity) showed increased activation of these neurons following sexual and aggressive behavioral encounters (Kabelik et al., 2013). While these studies found links between VP activity and behavior, it was not clear whether VP activity causally impacted behavioral expression, or whether neural input from the perception of and interaction with a conspecific may have led to the observed changes. The goal of the present study was thus to examine VP neuron activity in the absence of a conspecific, but within male green anoles whose stable boldness was established. Although not a test of causality, this approach allows us to ascertain the state of the vasopressinergic activity that likely precedes a behavioral encounter to predispose individuals toward greater or lesser behavioral expression once a conspecific in perceived. It should be noted, however, that the repeated testing to establish boldness could itself have long-lasting effects on VP chemoarchitecture and activity, much like the androgenic changes observed in the winner effect seen in some species (Fuxjager and Marler, 2010).

Experiments directly testing the causality of VP on behavior in lizards have also been conducted, though with inconclusive results (Campos et al., 2020; Dunham and Wilczynski, 2014). This may be partially due to logistic difficulties of targeted central injections, as these manipulations were via intraperitoneal injection. Thus, the resultant effects may be indirect due to peripheral binding of VP to smooth muscle, kidney, or pituitary receptors, rather than direct central manipulations. These green anole studies collectively found that VP manipulation decreased aggressive display to a mirror (though not to a conspecific), increased tongue flicking and chemical display, but also circulating glucocorticoid levels (the effects of which are then difficult to disentangle from those of VP itself). In another study, intraperitoneal administration of VP and the VP receptor antagonist Manning compound both failed to alter displayed aggression to a conspecific in male tree lizards (Kabelik, 2006).

Related to the present study, a recent green anole study examined gene expression in various brain regions of the five most bold and five most shy males from a distribution of fifty-seven animals (Kabelik et al., 2021). Interestingly, rather than regions containing VP neurons, the area containing the most expression differences between bold and shy males was the ventromedial hypothalamus. The androgen receptor was among the genes differentially expressed in this region, with increased expression in bold males. Testosterone, a ligand for these receptors, is known to regulate aggression-associated VP receptors in the ventrolateral hypothalamus (Delville et al., 1996). Androgens may also regulate VP receptors in the lateral ventromedial hypothalamus, a region both containing high densities of androgen receptors and showing aggressive display-inducted activation in lizards (Kabelik et al., 2008a; Rosen et al., 2002). Unfortunately, due to a lack of VP gene annotation in the green anole genome, the Kabelik et al. (2021) study could not determine whether nonapeptides including VP were differentially expressed between bold and shy males, though no differences in VP receptor expression in the ventromedial hypothalamus was detected.

### Ties to VP research in other vertebrate taxa

Apart from VP activity, the number of VP neurons present in a brain region also influences the amount of VP that can be released from that cell population. The number of VP neurons present in the bed nucleus of the stria terminalis of certain songbird species (as well as VP receptor densities in the lateral septum, a target site of these neurons) correlates positively with the degree of sociality (ranging from territorial to colonial) of the species (Goodson et al., 2006; Goodson and Wang, 2006). However, VP cells in the bed nucleus of the stria terminalis were barely detectable in a related brown anole study (Kabelik et al. 2013), and we were not able to discern any VP cells in that brain region in this green anole study. Vasopressin neurons of the PVN also play a related role in the regulation of social behaviors (Kelly and Goodson, 2014c) and knockdown of these neurons reduces gregariousness and alters displayed aggression in zebra finches (Kelly and Goodson, 2014a). Within the latter study, knockdown of VP in the PVN of male zebra finches caused increased aggression to opposite-sex individuals, while the same manipulation resulted in decreased aggression in females. While our study found no effects of neuron number within the PVN (or SON) on boldness, our results are nevertheless in line with those of Kelly and Goodson (2014a). Here, VP activity in the green anole PVN was correlated with decreased boldness in aggressive encounters (albeit toward a same-sex individual), which is consistent with those neurons reducing aggression in male zebra finches. However, the role of PVN VP neurons is complex, difficult to understand, and likely both sex- and species-specific. For instance, findings in male song sparrows find increased PVN VP activation following participation in a simulated agonistic encounter (Goodson and Evans, 2004), and aggression intensity is positively associated with PVN VP activity in male brown anoles (Kabelik et al., 2013). However, in goldfish, VP inhibits social approach when released within a hindbrain circuit (Thompson and Walton, 2004; Walton et al., 2010), which may be separate from forebrain circuitry regulating agonistic and anxiety related behaviors. This notion of separate cell groups exerting separate functions on social behavior is further supported in fish by work on African cichlids, where VP expression was found to be higher in gigantocellular cells of territorial than nonterritorial males, though a reverse finding was present in parvocellular neurons (Greenwood et al., 2008). Both of these cell groups are found within the preoptic area, an ancestral common region of VP production which gave rise to separate disparate populations in anamniotes (Goodson and Kabelik, 2009). In amniotes, both magnocellular and parvocellular VP neurons are present in the PVN (Kabelik et al., 2008c; Kawakami et al., 2021; Panzica et al., 1999), and this heterogeneity of cell types with separate functions may be one reason for the different functions and behavioral relationships attributed to VP neurons of the PVN across studies.

## Conclusions

Boldness of social interactions in male green anoles was found to be stable across weeks, as well as between agonistic and reproductive contexts. Baseline levels of VP activity (VP-pS6 colocalization) within both the PVN and SON were found to correlate inversely with the boldness of aggression toward males, though not with reproductive boldness toward females. Boldness was unrelated to measures of body size, circulating levels of glucocorticoids or sex steroids, or measures of VP cell number in the PVN and SON. The finding that VP activity correlates inversely with boldness suggests that VP modulates portions of the social decision-making network that regulate male aggression, and levels of VP release help determine individual variation in boldness during agonistic encounters. This hypothesis must be taken with caution, however, as a possible alternate hypothesis also exists, in that the extensive testing involved in determining measures of boldness could have produced long-lasting changes to VP neuronal activity. Further research will be required to help differentiate between these possibilities.

## Acknowledgements

LAO acknowledge that Stanford University resides on the ancestral and unceded land of the Muwekma Ohlone Tribe.

## Data Accessibility

Data from the analyses in this manuscript can be found at https://doi.org/10.6084/m9.figshare.16607780.

## References

Adriaenssens, B., Johnsson, J.I., 2011. Shy trout grow faster: exploring links between personality and fitness-related traits in the wild. Behav. Ecol. 22, 135–143. https://doi.org/10.1093/beheco/arq185

Albers, H.E., 2015. Species, sex and individual differences in the vasotocin/vasopressin system: Relationship to neurochemical signaling in the social behavior neural network. Front. Neuroendocrinol. https://doi.org/10.1016/j.yfrne.2014.07.001

Alfonso, S., Sadoul, B., Gesto, M., Joassard, L., Chatain, B., Geffroy, B., Bégout, M.-L., 2019. Coping styles in European sea bass: The link between boldness, stress response and neurogenesis. Physiol. Behav. 207, 76–85. https://doi.org/https://doi.org/10.1016/j.physbeh.2019.04.020

Ballew, N.G., Mittelbach, G.G., Scribner, K.T., 2017. Fitness consequences of boldness in juvenile and adult largemouth bass. Am. Nat. 189, 396–406. https://doi.org/10.1086/690909

Baugh, A.T., Schaper, S. V, Hau, M., Cockrem, J.F., de Goede, P., Oers, K. van, 2012. Corticosterone responses differ between lines of great tits (Parus major) selected for divergent personalities. Gen. Comp. Endocrinol. 175, 488–494. https://doi.org/https://doi.org/10.1016/j.ygcen.2011.12.012

Benjamini, Y., Hochberg, Y., 1995. Controlling the False Discovery Rate: A Practical and Powerful Approach to Multiple Testing. J. R. Stat. Soc. Ser. B 57, 289–300. https://doi.org/10.1111/j.2517-6161.1995.tb02031.x

Brown, C., Braithwaite, V.A., 2004. Size matters: a test of boldness in eight populations of the poeciliid Brachyraphis episcopi. Anim. Behav. 68, 1325–1329. https://doi.org/10.1016/j.anbehav.2004.04.004

Campos, S.M., Rojas, V., Wilczynski, W., 2020. Arginine vasotocin impacts chemosensory behavior during social interactions of Anolis carolinensis lizards. Horm. Behav. 124, 104772. https://doi.org/10.1016/j.yhbeh.2020.104772

Cao, R., Anderson, F.E., Jung, Y.J., Dziema, H., Obrietan, K., 2011. Circadian regulation of mammalian target of rapamycin signaling in the mouse suprachiasmatic nucleus. Neuroscience 181, 79–88. https://doi.org/10.1016/j.neuroscience.2011.03.005

Carter, C.S., 2017. The oxytocin-vasopressin pathway in the context of love and fear. Front. Endocrinol. (Lausanne). https://doi.org/10.3389/fendo.2017.00356

Colléter, M., Brown, C., 2011. Personality traits predict hierarchy rank in male rainbowfish social groups. Anim. Behav. 81, 1231–1237. https://doi.org/10.1016/j.anbehav.2011.03.011

Cornett, L.E., Kang, S.W., Kuenzel, W.J., 2013. A possible mechanism contributing to the synergistic action of vasotocin (VT) and corticotropin-releasing hormone (CRH) receptors on corticosterone release in birds. Gen. Comp. Endocrinol. 188, 46–53. https://doi.org/10.1016/j.ygcen.2013.02.032

Delville, Y., Mansour, K.M., Ferris, C.F., 1996. Testosterone facilitates aggression by modulating vasopressin receptors in the hypothalamus. Physiol. Behav. 60, 25–29. https://doi.org/10.1016/0031-9384(95)02246-5

Dunham, L.A., Wilczynski, W., 2014. Arginine vasotocin, steroid hormones and social behavior in the green anole lizard (Anolis carolinensis). J. Exp. Biol. 217, 3670–3676. https://doi.org/10.1242/jeb.107854

Félix, A.S., Cardoso, S.D., Roleira, A., Oliveira, R.F., 2020. Forebrain transcriptional response to transient changes in circulating androgens in a cichlid fish. G3 Genes, Genomes, Genet. 10, 1971–1982. https://doi.org/10.1534/g3.119.400947

Fuxjager, M.J., Marler, C.A., 2010. How and why the winner effect forms: Influences of contest environment and species differences. Behav. Ecol. 21, 37–45. https://doi.org/10.1093/beheco/arp148

Goodson, J.L., Evans, A.K., 2004. Neural responses to territorial challenge and nonsocial stress in male song sparrows: Segregation, integration, and modulation by a vasopressin V 1 antagonist. Horm. Behav. 46, 371–381. https://doi.org/10.1016/j.yhbeh.2004.02.008

Goodson, J.L., Evans, A.K., Wang, Y., 2006. Neuropeptide binding reflects convergent and divergent evolution in species-typical group sizes. Horm. Behav. 50, 223–236. https://doi.org/10.1016/j.yhbeh.2006.03.005

Goodson, J.L., Kabelik, D., 2009. Dynamic limbic networks and social diversity in vertebrates: From neural context to neuromodulatory patterning. Front. Neuroendocrinol. https://doi.org/10.1016/j.yfrne.2009.05.007

Goodson, J.L., Wang, Y., 2006. Valence-sensitive neurons exhibit divergent functional profiles in gregarious and asocial species. Proc. Natl. Acad. Sci. 103, 17013 LP – 17017. https://doi.org/10.1073/pnas.0606278103

Greenberg, N., Crews, D., 1990. Endocrine and behavioral responses to aggression and social dominance in the green anole lizard, Anolis carolinensis. Gen. Comp. Endocrinol. 77, 246– 255. https://doi.org/10.1016/0016-6480(90)90309-A

Greenwood, A.K., Wark, A.R., Fernald, R.D., Hofmann, H.A., 2008. Expression of arginine vasotocin in distinct preoptic regions is associated with dominant and subordinate behaviour in an African cichlid fish. Proc. R. Soc. B Biol. Sci. 275, 2393–2402. https://doi.org/10.1098/rspb.2008.0622

Hartline, J.T., Smith, A.N., Kabelik, D., 2017. Serotonergic activation during courtship and aggression in the brown anole, Anolis sagrei. PeerJ 2017, e3331. https://doi.org/10.7717/peerj.3331

Hattori, T., Wilczynski, W., 2009. Comparison of arginine vasotocin immunoreactivity differences in dominant and subordinate green anole lizards. Physiol. Behav. 96, 104–107. https://doi.org/10.1016/j.physbeh.2008.09.010

Kabelik, D., 2006. Neural mechanisms underlying the effects of testosterone on aggressive behavior in the tree lizard, Urosaurus ornatus. ProQuest Diss. Theses. Arizona State University, Ann Arbor.

Kabelik, D., Alix, V.C., Burford, E.R., Singh, L.J., 2013. Aggression- and sex-induced neural activity across vasotocin populations in the brown anole. Horm. Behav. 63, 437–446. https://doi.org/10.1016/j.yhbeh.2012.11.016

Kabelik, D., Alix, V.C., Singh, L.J., Johnson, A.L., Choudhury, S.C., Elbaum, C.C., Scott, M.R., 2014. Neural activity in catecholaminergic populations following sexual and aggressive interactions in the brown anole, Anolis sagrei. Brain Res. 1553, 41–58. https://doi.org/10.1016/j.brainres.2014.01.026

Kabelik, D., Crews, D., 2017. Hormones, Brain, and Behavior in Reptiles, in: Hormones, Brain and Behavior: Third Edition. pp. 171–213. https://doi.org/10.1016/B978-0-12-803592-4.00027-4

Kabelik, D., Crombie, T., Moore, M.C., 2008a. Aggression frequency and intensity, independent of testosterone levels, relate to neural activation within the dorsolateral subdivision of the ventromedial hypothalamus in the tree lizard Urosaurus ornatus. Horm. Behav. 54, 18–27. https://doi.org/10.1016/j.yhbeh.2007.09.022

Kabelik, D., Hofmann, H.A., 2018. Comparative neuroendocrinology: A call for more study of reptiles! Horm. Behav. 106, 189–192. https://doi.org/10.1016/j.yhbeh.2018.10.005

Kabelik, D., Julien, A.R., Ramirez, D., O’Connell, L.A., 2021. Social boldness correlates with brain gene expression in male green anoles. Horm. Behav. 133, 105007. https://doi.org/10.1016/j.yhbeh.2021.105007

Kabelik, D., Magruder, D.S., 2014. Involvement of different mesotocin (oxytocin homologue) populations in sexual and aggressive behaviours of the brown anole. Biol. Lett. 10, 20140566. https://doi.org/10.1098/rsbl.2014.0566

Kabelik, D., Weiss, S.L., Moore, M.C., 2008b. Steroid hormones alter neuroanatomy and aggression independently in the tree lizard. Physiol. Behav. 93, 492–501. https://doi.org/10.1016/j.physbeh.2007.10.008

Kabelik, D., Weiss, S.L., Moore, M.C., 2008c. Arginine vasotocin (AVT) immunoreactivity relates to testosterone but not territorial aggression in the tree lizard, Urosaurus ornatus. Brain. Behav. Evol. 72, 283–294. https://doi.org/10.1159/000174248

Kabelik, D., Weiss, S.L., Moore, M.C., 2006. Steroid hormone mediation of limbic brain plasticity and aggression in free-living tree lizards, Urosaurus ornatus. Horm. Behav. 49, 587–597. https://doi.org/10.1016/j.yhbeh.2005.12.004

Kabelik, D., Weitekamp, C.A., Choudhury, S.C., Hartline, J.T., Smith, A.N., Hofmann, H.A., 2018. Neural activity in the social decision-making network of the brown anole during reproductive and agonistic encounters. Horm. Behav. 106, 178–188. https://doi.org/10.1016/j.yhbeh.2018.06.013

Kanitz, E., Tuchscherer, M., Otten, W., Tuchscherer, A., Zebunke, M., Puppe, B., 2019. Coping style of pigs is associated with different behavioral, neurobiological and immune responses to stressful challenges. Front. Behav. Neurosci. https://doi.org/10.3389/fnbeh.2019.00173

Kawakami, N., Otubo, A., Maejima, S., Talukder, A.H., Satoh, K., Oti, T., Takanami, K., Ueda, Y., Itoi, K., Morris, J.F., Sakamoto, T., Sakamoto, H., 2021. Variation of pro-vasopressin processing in parvocellular and magnocellular neurons in the paraventricular nucleus of the hypothalamus: Evidence from the vasopressin-related glycopeptide copeptin. J. Comp. Neurol. 529, 1372–1390. https://doi.org/10.1002/cne.25026

Kelly, A.M., Goodson, J.L., 2014a. Hypothalamic oxytocin and vasopressin neurons exert sex-specific effects on pair bonding, gregariousness, and aggression in finches. Proc. Natl. Acad. Sci. U. S. A. 111, 6069–6074. https://doi.org/10.1073/pnas.1322554111

Kelly, A.M., Goodson, J.L., 2014b. Social functions of individual vasopressin-oxytocin cell groups in vertebrates: What do we really know? Front. Neuroendocrinol. 35, 512–529. https://doi.org/10.1016/j.yfrne.2014.04.005

Kelly, A.M., Goodson, J.L., 2014c. Personality is tightly coupled to vasopressin-oxytocin neuron activity in a gregarious finch. Front. Behav. Neurosci. https://doi.org/10.3389/fnbeh.2014.00055

Kelly, A.M., Kingsbury, M.A., Hoffbuhr, K., Schrock, S.E., Waxman, B., Kabelik, D., Thompson, R.R., Goodson, J.L., 2011. Vasotocin neurons and septal V1a-like receptors potently modulate songbird flocking and responses to novelty. Horm. Behav. 60, 12–21. https://doi.org/10.1016/j.yhbeh.2011.01.012

Ketterson, E.D., Nolan Val, J., 1999. Adaptation, exaptation, and constraint: A hormonal perspective. Am. Nat. 154, S4–S25. https://doi.org/10.1086/303280

Klingebiel, M., Dinekov, M., Köhler, C., 2017. Analysis of ribosomal protein S6 baseline phosphorylation and effect of tau pathology in the murine brain and human hippocampus. Brain Res. 1659, 121–135. https://doi.org/10.1016/j.brainres.2017.01.016

Knight, Z.A., Tan, K., Birsoy, K., Schmidt, S., Garrison, J.L., Wysocki, R.W., Emiliano, A., Ekstrand, M.I., Friedman, J.M., 2012. Molecular profiling of activated neurons by phosphorylated ribosome capture. Cell 151, 1126–1137. https://doi.org/10.1016/j.cell.2012.10.039

Koolhaas, J.M., de Boer, S.F., Coppens, C.M., Buwalda, B., 2010. Neuroendocrinology of coping styles: Towards understanding the biology of individual variation. Front. Neuroendocrinol. 31, 307–321. https://doi.org/10.1016/j.yfrne.2010.04.001

Korzan, W.J., Summers, C.H., 2004. Serotonergic response to social stress and artificial social sign stimuli during paired interactions between male Anolis carolinensis. Neuroscience 123, 835–845. https://doi.org/10.1016/j.neuroscience.2003.11.005

Korzan, W.J., Summers, T.R., Ronan, P.J., Renner, K.J., Summers, C.H., 2001. The role of monoaminergic nuclei during aggression and sympathetic social signaling. Brain. Behav. Evol. 57, 317–327. https://doi.org/10.1159/000047250

Larson, E.T., Summers, C.H., 2001. Serotonin reverses dominant social status. Behav. Brain Res. 121, 95–102. https://doi.org/10.1016/S0166-4328(00)00393-4

Lovern, M.B., Holmes, M.M., Wade, J., 2004. The green anole (Anolis carolinensis): A reptilian model for laboratory studies of reproductive morphology and behavior. ILAR J. https://doi.org/10.1093/ilar.45.1.54

Mayer, M., Shine, R., Brown, G.P., 2016. Bigger babies are bolder: effects of body size on personality of hatchling snakes. Behaviour 153, 313–323. https://doi.org/10.1163/1568539X-00003343

Mont, C., Hernandez-Pliego, P., Cañete, T., Oliveras, I., Río-Álamos, C., Blázquez, G., López-Aumatell, R., Martínez-Membrives, E., Tobeña, A., Flint, J., Fernández-Teruel, A., Mott, R., 2018. Coping-style behavior identified by a survey of parent-of-origin effects in the rat. G3 Genes, Genomes, Genet. 8, 3283–3291. https://doi.org/10.1534/g3.118.200489

Newman, S.W., 1999. The medial extended amygdala in male reproductive behavior. A node in the mammalian social behavior network. Ann. N. Y. Acad. Sci. 877, 242–257. https://doi.org/10.1111/j.1749-6632.1999.tb09271.x

O’Connell, L.A., Hofmann, H.A., 2012. Evolution of a vertebrate social decision-making network. Science (80-.). 336, 1154–1157. https://doi.org/10.1126/science.1218889

O’Connell, L.A., Hofmann, H.A., 2011. The vertebrate mesolimbic reward system and social behavior network: A comparative synthesis. J. Comp. Neurol. 519, 3599–3639. https://doi.org/10.1002/cne.22735

Panzica, G.C., Plumari, L., García-Ojeda, E., Deviche, P., 1999. Central vasotocin-immunoreactive system in a male passerine bird (Junco hyemalis). J. Comp. Neurol. 409, 105–117. https://doi.org/10.1002/(SICI)1096-9861(19990621)409:1<105::AID-CNE8>3.0.CO;2-8

Putman, B.J., Azure, K.R., Swierk, L., 2019. Dewlap size in male water anoles associates with consistent inter-individual variation in boldness. Curr. Zool. 65, 189–195. https://doi.org/10.1093/cz/zoy041

Qu, J., Fletcher, Q.E., Réale, D., Li, W., Zhang, Y., 2018. Independence between coping style and stress reactivity in plateau pika. Physiol. Behav. 197, 1–8. https://doi.org/10.1016/j.physbeh.2018.09.007

Réale, D., Dingemanse, N.J., Kazem, A.J.N., Wright, J., 2010. Evolutionary and ecological approaches to the study of personality. Philos. Trans. R. Soc. B Biol. Sci. 365, 3937–3946. https://doi.org/10.1098/rstb.2010.0222

Reaney, L.T., Backwell, P.R.Y., 2007. Risk-taking behavior predicts aggression and mating success in a fiddler crab. Behav. Ecol. 18, 521–525. https://doi.org/10.1093/beheco/arm014

Rosen, G., O’Bryant, E., Matthews, J., Zacharewski, T., Wade, J., 2002. Distribution of androgen receptor mRNA expression and immunoreactivity in the brain of the green anole lizard. J. Neuroendocrinol. 14, 19–28. https://doi.org/10.1046/j.0007-1331.2001.00735.x

Scherer, U., Kuhnhardt, M., Schuett, W., 2017. Different or alike? Female rainbow kribs choose males of similar consistency and dissimilar level of boldness. Anim. Behav. 128, 117–124. https://doi.org/10.1016/j.anbehav.2017.04.007

Sih, A., Bell, A., Johnson, J.C., 2004. Behavioral syndromes: an ecological and evolutionary overview. Trends Ecol. Evol. 19, 372–378. https://doi.org/10.1016/j.tree.2004.04.009

Sinervo, B., Miles, D.B., Frankino, W.A., Klukowski, M., DeNardo, D.F., 2000. Testosterone, endurance, and Darwinian fitness: Natural and sexual selection on the physiological bases of alternative male behaviors in side-blotched lizards. Horm. Behav. 38, 222–233. https://doi.org/10.1006/hbeh.2000.1622

Sluyter, F., Korte, S.M., Bohus, B., Van Oortmerssen, G.A., 1996. Behavioral stress response of genetically selected aggressive and nonaggressive wild house mice in the shock-probe/defensive burying test. Pharmacol. Biochem. Behav. 54, 113–116. https://doi.org/10.1016/0091-3057(95)02164-7

Smith, A.N., Kabelik, D., 2017. The effects of dopamine receptor 1 and 2 agonists and antagonists on sexual and aggressive behaviors in male green anoles. PLoS One 12. https://doi.org/10.1371/journal.pone.0172041

Smith, B.R., Blumstein, D.T., 2010. Behavioral types as predictors of survival in Trinidadian guppies (Poecilia reticulata). Behav. Ecol. 21, 919–926. https://doi.org/10.1093/beheco/arq084

Taborsky, M., Hofmann, H.A., Beery, A.K., Blumstein, D.T., Hayes, L.D., Lacey, E.A., Martins, E.P., Phelps, S.M., Solomon, N.G., Rubenstein, D.R., 2015. Taxon matters: promoting integrative studies of social behavior: NESCent Working Group on Integrative Models of Vertebrate Sociality: Evolution, Mechanisms, and Emergent Properties. Trends Neurosci. 38, 189–191. https://doi.org/10.1016/J.TINS.2015.01.004

Terranova, J.I., Ferris, C.F., Albers, H.E., 2017. Sex differences in the regulation of offensive aggression and dominance by Arginine-vasopressin. Front. Endocrinol. (Lausanne). https://doi.org/10.3389/fendo.2017.00308

Thompson, R.R., Walton, J.C., 2004. Peptide effects on social behavior: Effects of vasotocin and isotocin on social approach behavior in male goldfish (Carassius auratus). Behav. Neurosci. https://doi.org/10.1037/0735-7044.118.3.620

Thörnqvist, P.-O., McCarrick, S., Ericsson, M., Roman, E., Winberg, S., 2019. Bold zebrafish (Danio rerio) express higher levels of delta opioid and dopamine D2 receptors in the brain compared to shy fish. Behav. Brain Res. 359, 927–934. https://doi.org/10.1016/j.bbr.2018.06.017

Tokarz, R.R., 1995. Importance of androgens in male territorial acquisition in the lizard Anolis sagrei: an experimental test. Anim. Behav. 49, 661–669. https://doi.org/10.1016/0003-3472(95)80199-5

Tudorache, C., Slabbekoorn, H., Robbers, Y., Hin, E., Meijer, J.H., Spaink, H.P., Schaaf, M.J.M., 2018. Biological clock function is linked to proactive and reactive personality types. BMC Biol. 16, 148. https://doi.org/10.1186/s12915-018-0618-0

Veenema, A.H., Beiderbeck, D.I., Lukas, M., Neumann, I.D., 2010. Distinct correlations of vasopressin release within the lateral septum and the bed nucleus of the stria terminalis with the display of intermale aggression. Horm. Behav. 58, 273–281. https://doi.org/10.1016/j.yhbeh.2010.03.006

Veenema, A.H., Koolhaas, J.M., De Kloet, E.R., 2004. Basal and stress-induced differences in HPA axis, 5-HT responsiveness, and hippocampal cell proliferation in two mouse lines. Ann. N. Y. Acad. Sci. 1018, 255–265. https://doi.org/10.1196/annals.1296.030

Veenema, A.H., Meijer, O.C., De Kloet, E.R., Koolhaas, J.M., Bohus, B.G., 2003. Differences in basal and stress-induced HPA regulation of wild house mice selected for high and low aggression. Horm. Behav. 43, 197–204. https://doi.org/10.1016/S0018-506X(02)00013-2

Walton, J.C., Waxman, B., Hoffbuhr, K., Kennedy, M., Beth, E., Scangos, J., Thompson, R.R., 2010. Behavioral effects of hindbrain vasotocin in goldfish are seasonally variable but not sexually dimorphic. Neuropharmacology 58, 126–134. https://doi.org/10.1016/j.neuropharm.2009.07.018

Watt, M.J., Forster, G.L., Korzan, W.J., Renner, K.J., Summers, C.H., 2007. Rapid neuroendocrine responses evoked at the onset of social challenge. Physiol. Behav. 90, 567–575. https://doi.org/10.1016/j.physbeh.2006.11.006

Weiss, S.L., Moore, M.C., 2004. Activation of aggressive behavior by progesterone and testosterone in male tree lizards, Urosaurus ornatus. Gen. Comp. Endocrinol. 136, 282– 288. https://doi.org/10.1016/j.ygcen.2004.01.001

Wilczynski, W., Quispe, M., Muñoz, M.I., Penna, M., 2017. Arginine vasotocin, the Social Neuropeptide of Amphibians and Reptiles. Front. Endocrinol. (Lausanne). 8, 186. https://doi.org/10.3389/fendo.2017.00186

Woolley, S.C., Sakata, J.T., Crews, D., 2004a. Tyrosine hydroxylase expression is affected by sexual vigor and social environment in male Cnemidophorus inornatus. J. Comp. Neurol. 476, 429–439. https://doi.org/10.1002/cne.20236

Woolley, S.C., Sakata, J.T., Crews, D., 2004b. Evolutionary insights into the regulation of courtship behavior in male amphibians and reptiles. Physiol. Behav. 83, 347–360. https://doi.org/10.1016/j.physbeh.2004.08.021

Woolley, S.C., Sakata, J.T., Gupta, A., Crews, D., 2001. Evolutionary changes in dopaminergic modulation of courtship behavior in Cnemidophorus whiptail lizards. Horm. Behav. 40, 483–489. https://doi.org/10.1006/hbeh.2001.1713

Young, L.J., Greenberg, N., Crews, D., 1991. The effects of progesterone on sexual behavior in male green anole lizards (Anolis carolinensis). Horm. Behav. 25, 477–488. https://doi.org/10.1016/0018-506X(91)90015-A

