## Supplementary Data for "Aggressive but not reproductive boldness in male green anole lizards correlates with baseline vasopressin activity"

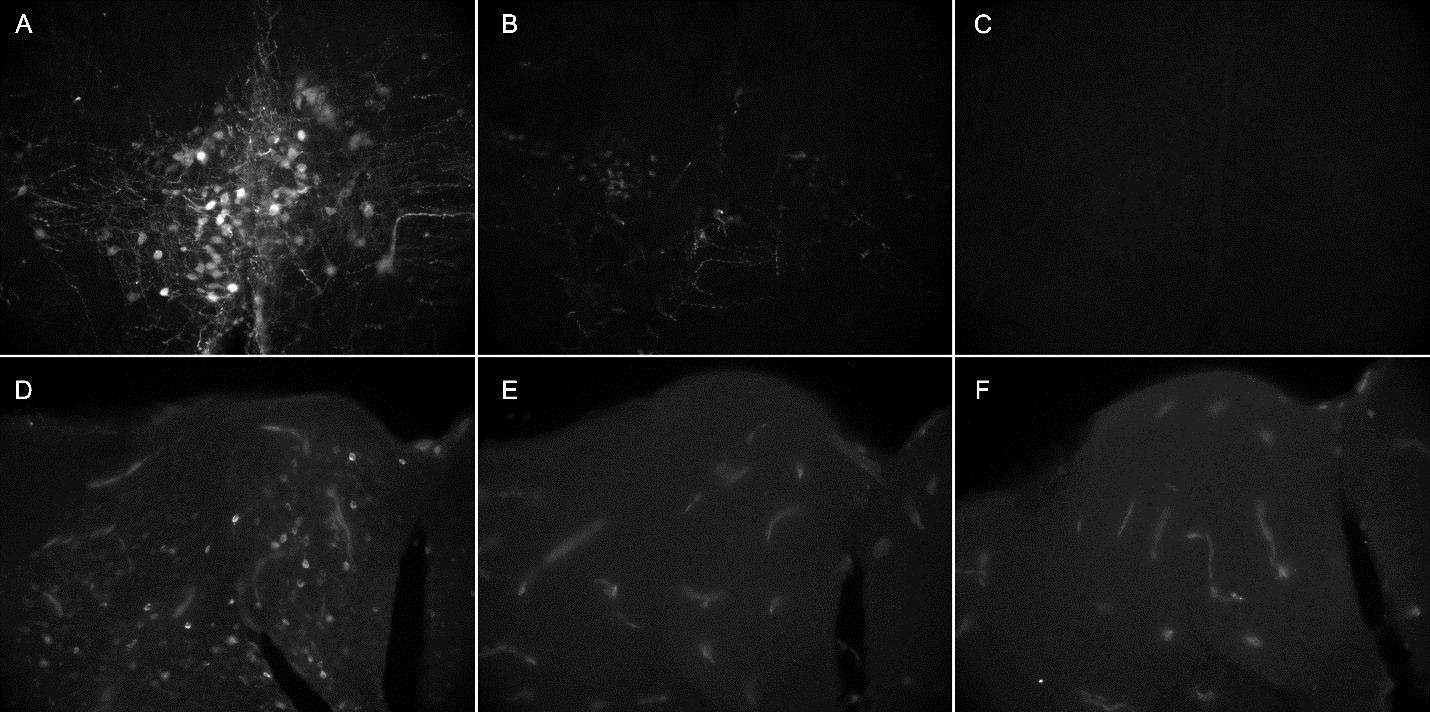

**Supplementary Figure 1.** **Preadsorption of VP and pS6.** Preadsorbing VP antibody with 50 µg/ml of peptide reduced signal (A), while 625 µg/ml of peptide completely eliminated signal (B). Preadsorbing pS6 antibody with 0.8 µg/ml of peptide (C) as well as 8 µg/ml of peptide (D) completely eliminated signal.

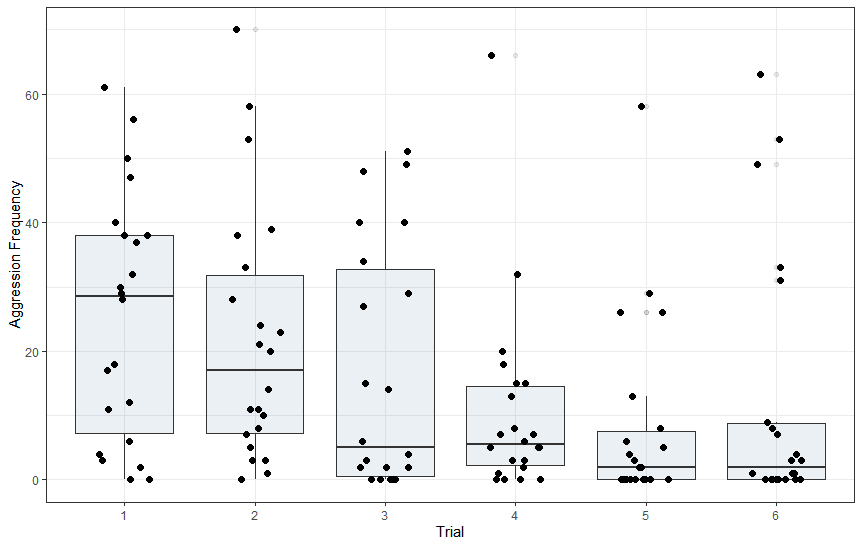

**Supplementary Figure 2:** Boxplots of displayed agression frequency across trials. Trials 1-3 took place on alternating days during week 1. Trials 4-6 were conducted with the same stimulus animals as trials 1-3 respectively, and took place on alternating days of week 2. Aggression frequency differed across trials (repeated-measures ANOVA, Greenhouse-Geisser, F=5.102, df=3.445, 72.346, p=0.002).

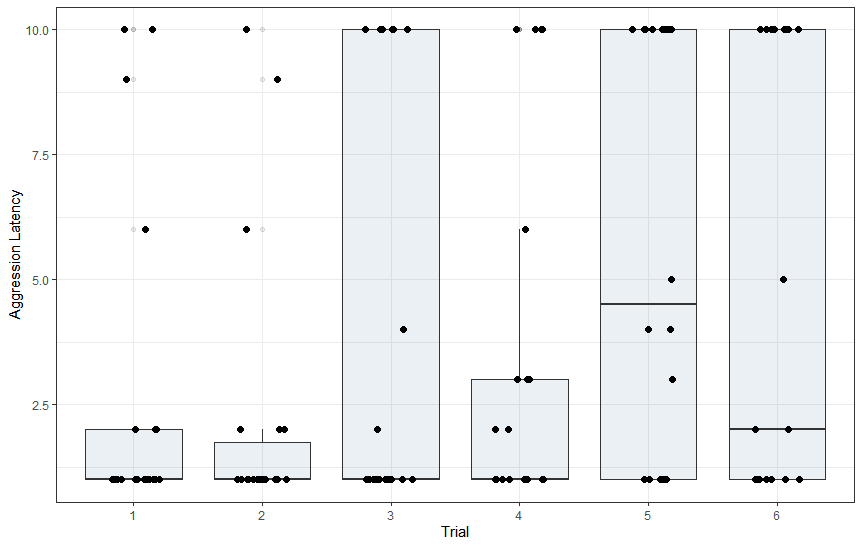

**Supplementary Figure 3:** Boxplots of displayed agression latency across trials. Trials 1-3 took place on alternating days during week 1. Trials 4-6 were conducted with the same stimulus animals as trials 1-3 respectively, and took place on alternating days of week 2. Aggression latency differed across trials (repeated-measures ANOVA, Greenhouse-Geisser, F=4.148, df=3.459, 72.631, p=0.006).

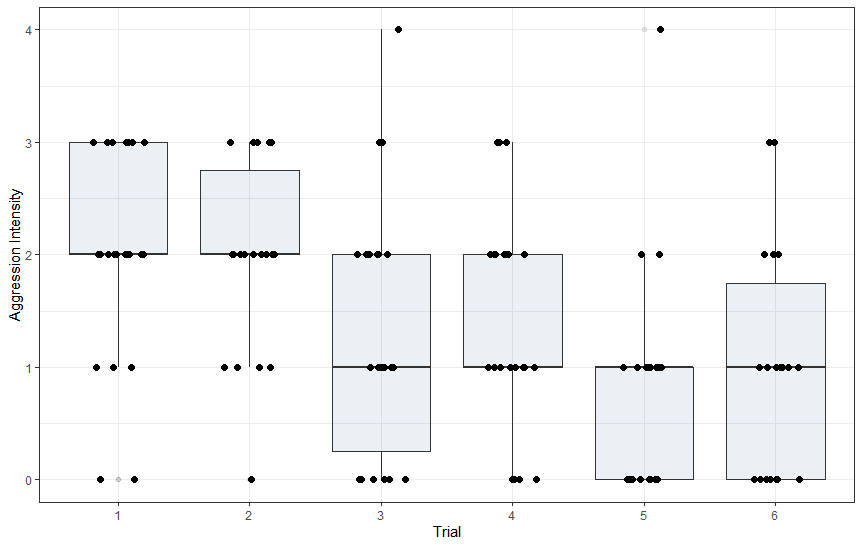

**Supplementary Figure 4:** Boxplots of displayed aggression intensity across trials. Trials 1-3 took place on alternating days during week 1. Trials 4-6 were conducted with the same stimulus animals as trials 1-3 respectively, and took place on alternating days of week 2. Aggression intensity differed across trials (Friedman test, χ^2^=28.207, N=22, p<0.001).

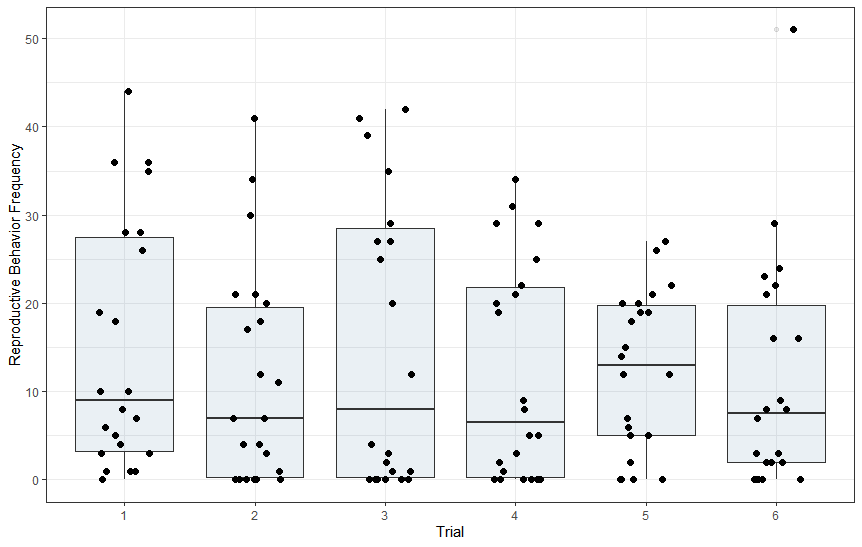

**Supplementary Figure 5:** Boxplots of displayed reproductive behavior frequency across trials. Trials 1-3 took place on alternating days during week 1. Trials 4-6 were conducted with the same stimulus animals as trials 1-3 respectively, and took place on alternating days of week 2. Reproductive behavior frequency did not differ across trials (repeated-measures ANOVA, Sphericity Assumed, F=0.917, df=5, 105, p=0.473).

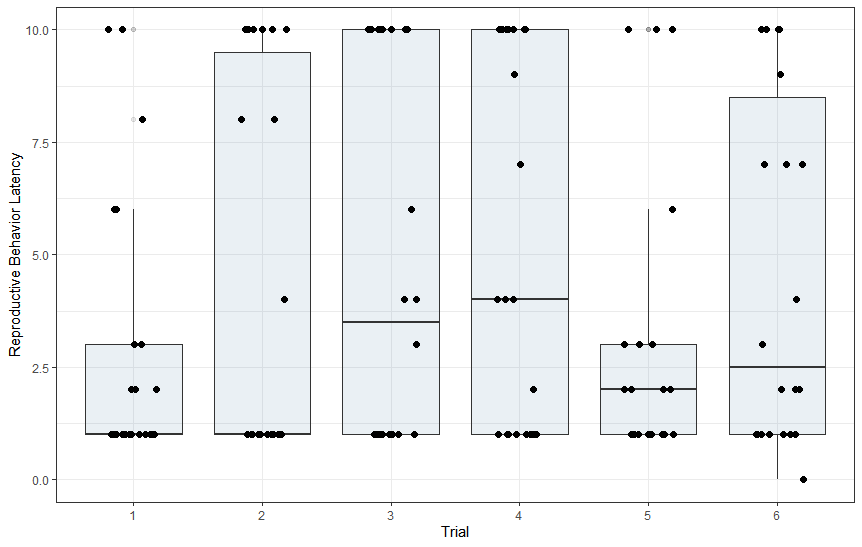

**Supplementary Figure 6:** Boxplots of displayed reproductive behavior latency across trials. Trials 1-3 took place on alternating days during week 1. Trials 4-6 were conducted with the same stimulus animals as trials 1-3 respectively, and took place on alternating days of week 2. Reproductive behavior latency did not differ across trials (repeated-measures ANOVA, Sphericity Assumed, F=1.723, df=5, 105, p=0.136).

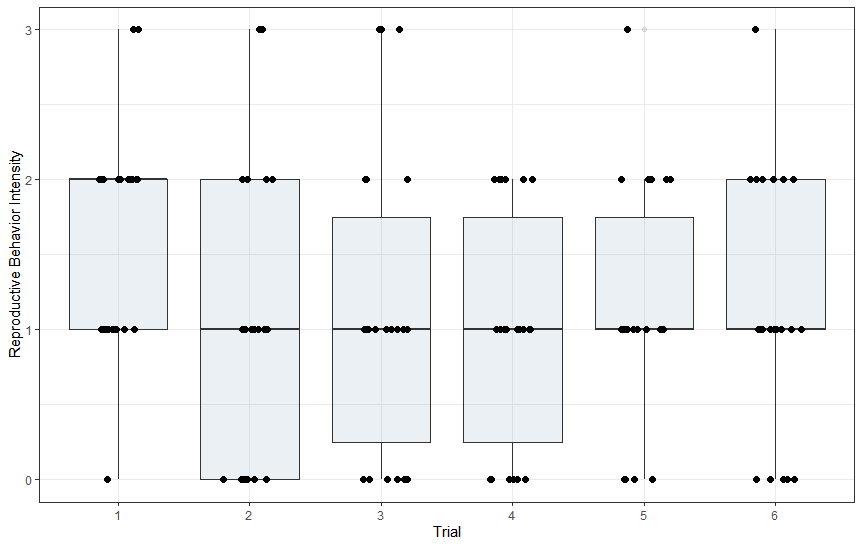

**Supplementary Figure 7:** Boxplots of displayed reproductive behavior intensity across trials. Trials 1-3 took place on alternating days during week 1. Trials 4-6 were conducted with the same stimulus animals as trials 1-3 respectively, and took place on alternating days of week 2. Reproductive behavior intensity did not differ across trials (Friedman test, χ^2^=10.342, N=22, p<0.066).
